# Estimating the efficacy of transcranial magnetic stimulation in small animals

**DOI:** 10.1101/058271

**Authors:** Lari M. Koponen, Jaakko O. Nieminen, Risto J. Ilmoniemi

## Abstract

The efficacy of transcranial magnetic stimulation (TMS) is determined by the magnitude and direction of the induced electric field in the cortex. The electric field distribution is influenced by the conductivity structure, in particular, the size of the head and the shapes of conductivity boundaries. We show that neglecting the head size can result in overestimating the stimulus intensity by a factor of 5–8 in the case of the rat brain. In the current modelling literature, the TMS-induced electric field is estimated with detailed computational simulations; however, in many experimental studies, less attention is paid on modelling. We attempt to bridge this gap by suggesting the use of simple simulations, for example with the spherical head model, when studying bioelectromagnetic phenomena.

## To the Editors of eLife

Murphy *et al.* (Murphy *et al.* 2016) showed in a rat model that transcranial magnetic stimulation (TMS) may cause neuronal inhibition. In this interesting study, TMS was delivered with a 70-mm figure-of-eight coil at a distance of 20–30 mm from the brain, which is typical in human experiments.

The stimulus intensity of 80–100% of the maximum stimulator output was estimated to induce an electric field in the rat cortex of approximately 150–200 V/m. This estimate was based on a study (Cohen et al., 1990) that reports induced electric field distributions in an infinite homogeneous medium for different coil and stimulator models. However, the electric field distribution due to TMS in a real head is strongly influenced by conductivity boundaries, and, as we will show, this contribution can be highly significant when the conductive volume is small compared to the dimensions of the coil.

The spherical head model (Sarvas, 1987) takes into account the effect of conductivity boundaries when estimating the induced electric field. Although both rat and human heads differ from perfect spherical symmetry, this model is reasonably accurate for TMS (Nummenmaa et al., 2013). We used the spherical model to estimate the electric field in the experimental condition of Murphy *et al.* and in some related cases. We modelled the magnetic field of the Magstim 70-mm figure-of-eight coil (Thielscher and Kammer, 2002) assuming maximum output of the Magstim Rapid^2^ stimulator for single-pulse stimulation with the same coil (Nieminen et al., 2015), as used by Murphy et al. (personal communication with Murphy, 18^th^ May 2016), and applied the reciprocity theorem to obtain the intracranial electric field (Heller and van Hulsteyn, 1992). The computational model is described in more detail in the methods section.

In (Figure 1), we show the induced electric field distributions in human and rat cortices, assuming head radii of 85 and 15 mm and scalp-to-cortex distances of 15 and 3 mm, respectively. With identical coil-to-cortex separation, the electric field in the rat cortex was found to be just 32% of that in the human cortex. If the coil was placed against the rat scalp, the maximum electric field in the rat cortex would still be only 59% of that in the human cortex.

**Figure 1:**
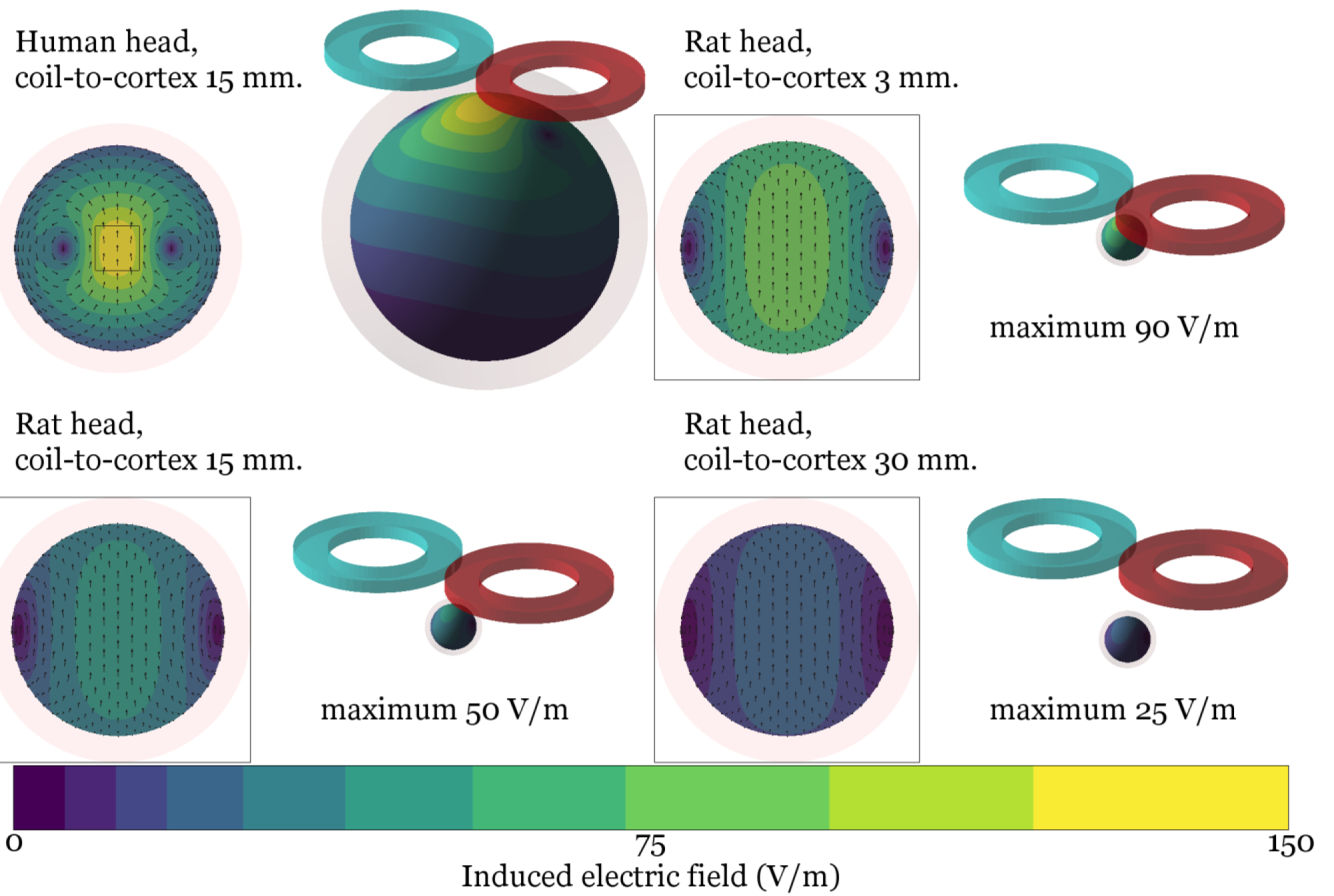
The TMS-induced electric field for a 70-mm figure-of-eight coil in various head geometries at 100– maximum stimulator output of Magstim Rapid^2^. The dimensions of the squares are 30 by 30 mm.

Our analysis suggests that the stimuli of Murphy *et al.* could have been below the motor threshold, thus explaining why they saw no behavioural responses in their rats. Researchers should be cautious when extrapolating the induced electric field values from one geometry to another. The differences in the electric field intensities highlight the importance of proper calibration (Nieminen et al., 2015) combined with adequate simulations when studying bioelectromagnetic phenomena.

## Methods

We utilised the reciprocity theorem to compute the TMS-induced electric field. In TMS, a changing current in the TMS coil windings induces an electric field distribution in the brain, whereas in magnetoencephalography (MEG) a source current distribution in the brain produces magnetic flux passing through the pickup coil windings. According to the reciprocity theorem, the ratio between the induced electric field at any point in any direction (in volts per metre) and the rate of change in the coil current (in amperes per second) in TMS equals minus one times the ratio between the coil flux (in weber) and the source current magnitude at the same point in the same direction (in ampere-metres) in MEG. The reciprocity theorem results from Maxwell’s equations in the low-frequency regime, and, as shown by Heller and van Hulsteyn (Heller and van Hulsteyn, 1992), is applicable to the frequencies present in a TMS pulse.

Thanks to reciprocity, we could use the Sarvas formula (Sarvas, 1987) to compute the induced electric field in the spherical model (Table 1); this was implemented with Matlab (R2015b, https://www.mathworks.com). We validated the resulting computational model for TMS-induced electric-field lead-field matrix by comparing its results to independently obtained induced-electric-field data from (Koponen et al., 2015). The implementation is available from the corresponding author for academic use.

**Table 1:**
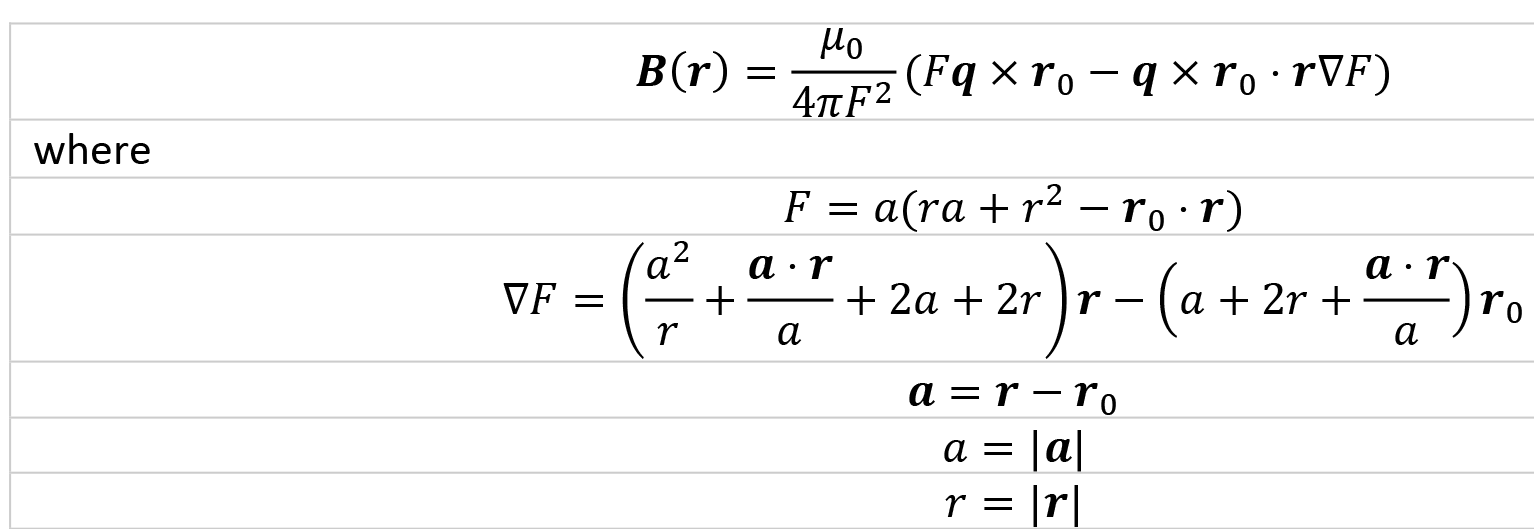
The magnetic field ***B*** at a location ***r*** outside a spherically symmetric conducting volume, centred at the origin, due to a source current dipole ***q*** at a location ***r***_0_ within the volume. *μ_0_* is the permeability of free space.

The TMS-coil model was built from a set of magnetic dipoles similarly to (Thielscher and Kammer, 2002) using the geometry of the Magstim 70-mm figure-of-eight coil from the same article, and the maximum stimulator output of the Magstim Rapid^2^ system from (Nieminen et al., 2015). We used the resulting computational model to study the TMS-induced electric field in several conditions.

## Acknowledgements

This work was funded by the Finnish Cultural Foundation and the Academy of Finland (Decisions No. 255347 and 265680).

## References

Cohen LG, Roth BJ, Nilsson J, Dang N, Panizza M, Bandinelli S, Friauf W, Hallett M. 1990. Effects of coil design on delivery of focal magnetic stimulation. Technical considerations. Electroencephalography and Clinical Neurophysiology 75:350–57. doi: 10.1016/0013-4694(90)90113-X

Heller L, van Hulsteyn DB. 1992. Brain stimulation using electromagnetic sources: theoretical aspects. Biophysical Journal 63:129–38. doi: 10.1016/S0006-3495(92)81587-4

Koponen LM, Nieminen JO, Ilmoniemi RJ. 2015. Minimum-energy coils for transcranial magnetic stimulation: application to focal stimulation. Brain Stimulation 8:124–34. doi: 10.1016/j.brs.2014.10.002

Murphy SC, Palmer, LM, Nyffeler T, Muri RM, Larkum ME. 2016. Transcranial magnetic stimulation (TMS) inhibits cortical dendrites. eLife 5:e13598. doi: 10.7554/eLife.13598

Nieminen JO, Koponen LM, Ilmoniemi RJ. 2015. Experimental characterization of the electric field distribution induced by TMS devices. Brain Stimulation 8:582–89. doi: 10.1016/j.brs.2015.01.004

Nummenmaa A, Stenroos M, Ilmoniemi RJ, Okada YC, Hamalainen MS, Raij T. 2013. Comparison of spherical and realistically shaped boundary element head models for transcranial magnetic stimulation. Clinical Neurophysiology 124:1995–2007. doi: 10.1016/j.clinph.2013.04.019

Sarvas J. 1987. Basic mathematical and electromagnetic concepts of the biomagnetic inverse problem. Physics in Medicine and Biology 32:11–22. doi: 10.1088/0031-9155/32/1/004

Thielscher A, Kammer, T. 2002. Linking physics with physiology in TMS: a sphere field model to determine the cortical stimulation site in TMS. NeuroImage 17:1117–30. doi: 10.1006/nimg.2002.1282

